# Generation of two Multipotent Mesenchymal Progenitor Cell Lines Capable of Osteogenic, Mature Osteocyte, Adipogenic, and Chondrogenic Differentiation

**DOI:** 10.1101/2020.11.19.385138

**Authors:** Matthew Prideaux, Christian S. Wright, Megan L. Noonan, Xin Yi, Erica L. Clinkenbeard, Elsa Mevel, Jonathan A. Wheeler, Sharon Byers, Uma Sankar, Kenneth E. White, Gerald J. Atkins, William R. Thompson

## Abstract

Differentiation of multi-potent mesenchymal progenitor cells give rise to several tissue types including bone, cartilage, and adipose. In addition to the complication arising from the numerous spatial, temporal, and hormonal factors that regulate lineage allocation, targeting of these cells *in vivo* is challenging, making mesenchymal progenitor cell lines valuable tools to study both tissue development and the differentiated cell types. Mesenchymal stem cells (MSCs) can be isolated from humans and animals; however, obtaining homogenous, responsive cells in a reproducible fashion can be problematic. As such, we have developed two novel mesenchymal progenitor cell (MPC) lines, MPC1 and MPC2, which were generated from the bone marrow of male C57BL/6 mice. These cells were immortalized using the temperature sensitive large T-antigen, allowing for thermal control of proliferation and differentiation. Both MPC1 and MPC2 cell lines are capable of osteogenic, adipogenic, and chondrogenic differentiation. Under osteogenic conditions both cell lines formed discrete mineralized nodules, staining for alizarin red and alkaline phosphatase, while expressing high levels of osteogenic genes including *Sost*, *Fgf23*, and *Dmp1*. *Sost* and *Dmp1* mRNA levels were drastically reduced with parathyroid hormone, thus recapitulating *in vivo* responses. MPC cells secreted both the intact (iFGF23) and *C*-terminal (cFGF23) forms of endocrine hormone FGF23, which was upregulated in the presence of 1,25 dihydroxy vitamin D (1,25D). In addition to osteogenic differentiation, both cell lines also rapidly entered the adipogenic lineage, expressing several adipose markers after only 4 days in adipogenic media. MPC cells were also capable of chondrogenic differentiation, displaying increased expression of common cartilage genes including aggrecan, sox9, and cartilage oligomeric matrix protein. With the ability to differentiate into multiple mesenchymal lineages and mimic in vivo responses of key regulatory genes/proteins, MPC cells are a valuable model to study factors that regulate mesenchymal lineage allocation as well as the mechanisms that dictate transcription, protein modification, and secretion of these factors.

## Introduction

The ability of mesenchymal stem cells (MSCs) to differentiate into multiple lineages makes them a valuable tool for the investigation of tissue development and responses to various stimuli. While MSCs are present in numerous tissues, bone marrow derived MSCs are essential for cartilage formation, bone remodeling and repair, and can also form into adipose tissue [1]. As the bone marrow niche contains numerous differentiated and progenitor cell types, studying the properties of MSCs *in vivo* is challenging. Thus, cultured MSCs are useful to determine the cell-specific responses of MSCs apart from other cell types; however, there are several challenges with current cell models.

Primary MSCs can be extracted from various tissues, but are most commonly isolated from adipose and bone marrow [2] in both human and animal models. Although culture conditions can be optimized to promote growth of MSCs over other cell types, these cultures typically remain highly heterogenous, containing hematopoietic cells and other cell types. In addition to the contamination of other cell types, there is wide variation among donors (or animals) and even between isolations within the same donor. Primary MSC cultures often display differing growth kinetics and variation in the proportion of cell populations therein. Such variabilities create challenges in obtaining consistent phenotypic and functional results and may even lead to incorrect interpretation of data [2].

In contrast to primary cells, immortalized cell lines provide greater homogeneity and are thus capable of producing more consistent experimental outcomes. Numerous human MSC cell lines are available through commercial vendors, some of which are immortalized, enabling greater expansion in the laboratory setting. Human MSC cells derive from various tissues, most commonly adipose aspirates, but also from bone marrow [3]. While the use of human MSCs provides a powerful tool that may be more easily translated to human studies, human cells often vary greatly in their responses compared to cells derived from animals. Furthermore, most of the available MSC lines require complicated protocols to induce differentiation, often including the use of proprietary cell culture medium, which is both costly and introduces undefined components (likely various growth factors) that may influence downstream experimental outcomes. Additionally, the majority of these cell lines require upwards of 4 weeks in culture to achieve osteogenic and adipogenic differentiation [4–6].

Mice are an extremely useful biological model, and the ability to induce transgenic modifications in mice makes them a powerful tool for discovery. As such, mouse cell lines are useful for *in vitro* confirmation of *in vivo* results, often providing essential data to translate findings from animal to human studies. Unfortunately, there are relatively few established immortalized mouse MSC cell lines, and the lines that are available have similar challenges as human cell lines. In the absence of readily available MSC cell lines many groups isolate primary cells for each experiment, requiring the use of numerous mice. A few groups have developed improved methods for the isolation of MSCs from mice that allow for multiple passages of the isolated cells [7,8]. This is especially useful to generate cells from mice with transgenic modifications; however, the limitations with heterogeneity and contamination with other cell types remain.

To overcome some of the common issues with primary MSC cultures, we developed two novel multi-potent cell lines capable of differentiating into the osteogenic, adipogenic, and chondrogenic lineages. These cells were isolated from the bone marrow of C57BL/6 mice and are referred to as Murine Progenitor Cells 1 and 2 (MPC1 & MPC2). MPC1 and MPC2 cells were expanded from single cell clones and harbor the temperature sensitive large T-antigen, providing a unique ability to regulate proliferation and differentiation based on the incubation temperature. In addition to the tri-lineage capacity, MPC cells produce very high expression of several commonly studied molecules. In particular, both MPC1 and MPC2 cells express and secrete the intact (iFGF23) and *C*-terminal forms of fibroblast growth factor 23 protein (cFGF23). While several other cell lines produce *Fgf23* mRNA, there are very few available that secrete FGF23 protein. As such, these cells provide a useful tool not only to study responses of mesenchymal progenitors, but also to examine development of multiple tissue types.

## Materials and Methods

### Reagents

Cell culture media, trypsin-EDTA, and antibiotics were purchased from Invitrogen (Carlsbard, CA). Fetal bovine serum (FBS) was purchased from Atlanta Biologicals (Atlanta, GA). Ascorbic acid, β-glycerophosphate, alizarin red, and the alkaline phosphatase staining kit were obtained from Sigma Aldrich (St. Louis, MO).

### Antibodies

Antibodies (Abs) recognizing PPARγ (#2443), GAPDH (#5174), and perilipin (#9349) were purchased from Cell Signaling (Danvers, MA). Antibody against fatty acid binding protein 4 (FABP4) was purchased from ProSci, Inc. (Poway, CA) (XG6174). The anti-adiponectin Ab (PA1-054) was obtained from Affinity BioReagents (Rockford, IL). Aggrecan Ab was from Abcam (ab3778). Antibody against collagen I was purchased from Millipore (AB765P), and the collagen X Ab was purchased from Calbiochem (234196).

### Cell Isolation and Culture Conditions

Mesenchymal progenitor cells were isolated from 8-week-old male C57BL/6 mice as previously described[8]. Briefly, mice were euthanized by CO2 asphyxiation followed by cervical dislocation. Femurs and tibias were dissected and kept on ice in Roswell Park Memorial Institute (RPMI) media with FBS (9%, v/v), horse serum (9%, v/v), and penicillin/streptomycin (100 μg/ml). Marrow was flushed from long bones and passed through a nylon mesh cell strainer (70 μm). Cells were spun down (7,000 rpm), resuspended in RPMI media, and plated in a T175 flask. Cells were cultured for two passages in RPMI media, then passaged twice more in Iscove’s Modified Dulbecco’s Medium (IMDM) containing FBS (9%, v/v), horse serum (9%, v/v), and penicillin/streptomycin (100 μg/ml). Cells then were expanded and cryogenically preserved. All mouse handling and cell isolation was performed under protocols approved by the Indiana University Institutional Animal Care and Use Committee.

### Immortalization Plasmid

The temperature-sensitive large T antigen SV40 sequence, SVU19tsa58, was digested from pZipSVU19tsa58 plasmid (a gift from Parmjit Jat, UCL, UK) using the BamH1 restriction site. The sequence was ligated into the pLVX Puro lentiviral plasmid (Clontech, CA) using the same restriction sites and sequenced to confirm correct orientation. Plasmid DNA was amplified in One Shot Stbl3 E. coli (Thermo-Fisher Scientific) and purified using a Plasmid Maxiprep kit (Qiagen).

### Lentivirus Generation

Human embryonic kidney (HEK) 293 T cells were transiently transfected with the immortalizing plasmid and plasmids encoding for Tat, Rev, Gag/Pol and VSV-G using Fugene-6 transfection reagent as per the manufacturer’s instructions (Roche) for 8 hours. Media was replaced with DMEM containing FBS (10%, v/v) and cells were incubated for 48 h. The media containing virus particles was collected, centrifuged (2000 rpm) to pellet cell debris, the supernatant was passed through a filter (0.45 μm), and stored at −80°C.

### Lentiviral Transduction

MSCs were seeded at a density of 2×10^4^ cells/cm^2^ in T25 flasks the day prior to transduction. The following day, media was removed and replaced with a mixture of culture media (50%) and crude virus (50%) containing Polybrene (8 μg/ml, Sigma). Transduction media was removed and replaced with fresh culture media 24 h later. After an additional 48 h, fresh media containing puromycin (2 μg/ml, Sigma) was added to select for infected cells. Puromycin selection was maintained for 10 days.

### Transfection

MPC cells were plated in 6-well plates (100,000 cells/well). After 24 h cells were transfected with an eGFP vector (3 μg; Clontech) using Fugene-6 HD reagent according to the manufacturer's protocol. Cells were incubated at 33° C for the for at least 24 h before being imaged with a fluorescence on a microscope (Leica).

### Single Cell Cloning

Stably transfected cells were cloned by limiting dilution in 96-well culture plates. Over 20 clones were created, of which 10 were differentiated in osteogenic media containing alpha minimal essential media (α-MEM), FBS (10%, v/v), β-glyerophosphate (5mM) and ascorbic acid (50μg/ml) for 21 days. The two mesenchymal progenitor cell clones showing the greatest mineralization, as determined by alizarin red staining, were selected for further characterization and designated as “MPC1” and “MPC2”.

### Differentiation Conditions

Cells were cultured at 33°C to allow for proliferation. For osteogenic and adipogenic differentiation MPC1 and MPC2 cells were seeded at 6,000 - 10,000 cells/cm^2^ in six-well dishes (Corning, Corning, NY). Cells were incubated overnight (ON) in IMDM media to allow them to adhere. The following day cells were moved to 37°C and IMDM media was replaced with osteogenic media consisting of α-MEM ascorbic acid (50 μg/ml) and β-glycerophosphate (10 mM) or adipogenic media containing dexamethasone (0.1 μM), insulin (5 μg/ml) and indomethacin (50 μM). For osteogenic cultures, media was changed every 48 h. To determine osteocytic responses of endocrine factors MPC cells were differentiated in osteogenic media for 28 days followed by addition of PTH (50 mM) or 1,25D (10 nM) for 24 hours, after which time RNA or media were collected for analysis.

Induction of chondrogenic differentiation was accomplished by adding 250,000 – 500,000 MPC1 or MPC2 cells to polypropylene tubes containing growth media containing sodium L-ascorbate (50 nM), insulin (6.25 μg/mL), sodium selenite (6.25 ng/mL) and penicillin/streptomycin (1%, v/v) in DMEM or with chondrogenic media composed of insulin (6.25 μg/mL), transferrin (6.25 μg/mL), sodium selenite (6.25 ng/mL), sodium L-ascorbate (50 nM), dexamethasone (10^−8^ M), and TGF-β1 (10 ng/mL) in DMEM. Cells were centrifuged for 5 min (250xg), fitted with vented caps and incubated for 48 h in normoxia. After 48 h chondrogenic cultures were moved to hypoxic conditions (5% O_2_) and cultured for 28 days.

### Western Blotting

Whole cell lysates were prepared using radio immunoprecipitation assay (RIPA) lysis buffer (150 mM NaCl, 50 mM Tris HCl, 1 mM EGTA, 0.24% (w/v) sodium deoxycholate,1% (w/v) Igepal, pH 7.5) with protease and phosphatase inhibitors. Inhibitors including NaF (1 mM) and Na_3_VO_4_ (1 mM), aprotinin (1 μg/ml), leupeptin (1 μg/ml), pepstatin (1 μg/ml), and phenylmethylsulfonylfluoride (PMSF, 1 mM) were added fresh, just prior to lysis. Total protein lysates (20 μg) were separated on SDS polyacrylamide gradient gels (4-12%) and transferred to polyvinylidene difluoride (PVDF) membranes, as described previously[9]. Membranes were blocked with milk (5%, w/v) diluted in tris-buffered saline containing tween-20 (TBS-T, 0.01%, v/v). Blots then were incubated ON at 4°C with the appropriate primary antibody. Blots were washed and incubated with horseradish peroxidase-conjugated secondary antibody (1:5000 dilution) (Cell Signaling) at RT for one hour with chemiluminescent detection using ECL Plus substrate (Amersham Biosciences, Piscataway, NJ). Images were developed and acquired with an iBright CL1000 machine (Applied Biosystems).

### Real Time PCR

Total RNA was isolated by using the RNeasy kit (Qiagen, Germantown, MD) as described previously[10]. mRNA was reverse transcribed, and genes were amplified with a BioRad CFX Connect^TM^ qPCR machine, using gene-specific primers (Table 1), as previously described[11]. PCR products were normalized to *Gapdh* mRNA expression and quantified using the ΔΔCT method.

### Alizarin red and Alkaline phosphatase staining

Mineralization was induced on confluent monolayers in 12-well plates by addition of osteogenic media. Monolayers were washed with 1X PBS and fixed for 1 hour with cold 70% ethanol, then washed 3 times with excess dH_2_O prior to addition of 1 mL of 2% w/v Alizarin Red S (pH 4.2) per well. The plates were incubated in the dark at room temperature for 10 min. After removal of unincorporated dye, wells were washed three times with dH_2_O, reaspirated, and stored at room temperature. In identical cultures for the mineralization assay, monolayers of MPC1 and MPC2 cells were washed twice with 1X PBS and fixed for 30 minutes in 4% PFA. PFA was removed and cells were washed again in 1X PBS, then stained using the Leukocyte Alkaline Phosphatase Kit (Sigma) as previously described [12]. Plates were stored at room temperature. Images of wells were taken at 10X magnification on an inverted Leica microscope.

### FGF23 protein detection

To test FGF23 protein production, MPC1 or MPC2 cells were seeded on 6-well plates and grown to confluence. Cells were differentiated in osteogenic media for 14 or 21 days then treated with 10^−8^M 1,25(OH)_2_ vitamin D (Sigma) or vehicle (DMSO) for 24 hrs. The media was removed centrifuged to remove unattached cells and debris. Media was concentrated in Amicon Ultra Centrifugal Filters (Milipore) and stored at −80°C. The adherent cells were lysed with 300 μL of 1X Lysis buffer (Cell Signaling Technologies, Inc., Danvers, MA, USA) with 1 μg/mL 4-(2-aminoethyl) benzenesulfonyl fluoride hydrochloride (AEBSF) protease inhibitor (Sigma-Aldrich, Inc.) according to the manufacturer’s directions. Total cell lysate protein concentrations were determined with the Better Bradford Kit (Thermo-Fisher Scientific) according to the manufacturer’s instructions. Secreted FGF23 protein was assessed using both the rodent-specific ‘intact’ FGF23 (‘iFGF23’) and ‘*C*-terminal’ (or ‘total’) ‘cFGF23’ ELISAs (Quidel Laboratories, Inc.) and normalized to total protein concentration.

### Histological Processing and Immunohistochemistry

Chondrogenic pellets were fixed in neutral buffered formalin (10%, v/v) for 48 h and embedded in paraffin. Paraffin embedded pellets were sectioned (10 μm), deparaffinized, and rehydrated prior to staining with either Safranin-O/Fast Green or Alcian Blue, as previously described[13].

Immunohistochemical staining of chondrogenic pellets was performed by first deparaffinizing sections and rehydration of sections. Epitope retrieval was achieved by incubation with chondroitinase ABC (2 mg/ml) for 30 min at RT (aggrecan and collagen X) or by incubation in citrate buffer (10 mM, pH 6.0) for 20 min at 90°C, followed by a 30 min incubation at room temperature (RT). Sections were rinsed with distilled/deionized water (ddH_2_O) then inactivated by exposure to H_2_O_2_ for 20 min. at RT and then blocked in PBS containing BSA (1%, w/v) and goat serum (5%, v/v) for 30 min at RT. Sections were incubated with primary Ab (1:100) in a humidified chamber at 4°C ON. After rinsing with PBS, horse radish peroxidase-conjugated secondary Ab was added for 1 h at RT and rinsed again with PBS. Sections then were exposed to 3,3’ Diaminobenzidine (DAB) chromogenic substrate, rinsed with ddH_2_O, counterstained with Gill No. 1 hematoxylin, dehydrated, cleared with Xylene, and mounted on slides prior to imaging.

### Statistical Analyses

Data were expressed as means ± SE. Statistical significance was analyzed using Student’s t-test or two-way ANOVA, allowing for unequal variance (Prism GraphPad, LA Jolla, CA). All assays were replicated at least three times, using biological replicates, to assure reproducibility.

## Results

### Osteogenic Induction of MPC1 and MPC2 Cells form Mineralized Nodules

MPC1 and MPC2 cells were exposed to osteogenic media (OM) for 7-28 days, followed by alizarin red staining for calcium deposition. Compared to the maintenance growth media, when cultured in osteogenic media both MPC1 and MPC2 cells displayed similar alizarin red staining when cultured in osteogenic media (Fig. 1A, B). Following 7 days of osteogenic differentiation, alizarin red staining of MPC2 cells (Fig. 1B) appeared more extensive than that of MPC1 cells (Fig. 1A). Staining of MPC2 also appeared greater at the 14-day timepoint compared to MPC1; however, the staining abundance was indistinguishable at 21 and 28 days of differentiation, where mineralization of nearly the entire well was observed. While the alizarin red stain was more diffuse across the culture dish in cells exposed to differentiation media for 28 days, distinct mineralized nodules were apparent at 7, 14, and 21 days of differentiation (Fig. 1).

**Figure 1.**
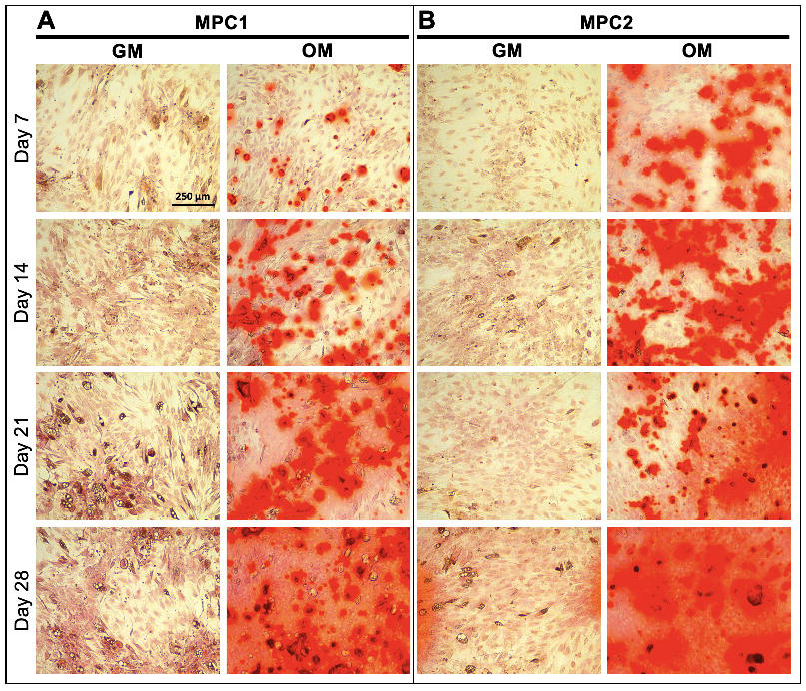
Time course of osteogenic mineralization for MPC1 and MPC2 cells. MPC1 (A) and MPC2 (B) cells were cultured in growth media (GM) or osteogenic media (OM) and stained with alizarin red to examine mineralization over the differentiation time course. (A) Mineralization of MPC1 cells cultured in OM began at day 7, which was strongly increased at Day 14, and even more robust at Day 21. (B) MPC2 cells at Day 7 in OM exhibited more robust staining compared to MPC1 at the same time. Alizarin red stain in OM continually increased with duration of differentiation. At day 28 of OM, both cell types developed similar and robust mineralization. In contrast to MPC1, MPC2 cells exhibited modest mineralization at Day 28 when maintained in the growth media. Each line was tested in triplicate. Images were captured using a 10X magnification lens.

MPC cells exposed to osteogenic differentiation conditions were also stained for the osteoblast marker alkaline phosphatase (Alkphos). The pattern of Alkphos staining was similar to that of alizarin red. Following 7 days of osteogenic differentiation MPC1 and MPC2 cells showed minimal Alkphos staining (Fig. S1A, S1B). At day 14, Alkphos staining was more apparent in cells exposed to osteogenic media in both MPC1 and MPC2 cells compared to growth media. Staining for Alkphos was increased in MPC1 and MPC2 cells exposed to osteogenic media at day 21 compared to cells cultured in growth media. Staining was approximately equal for MPC1 and MPC2 cells at day 21. After 28 days in culture, MPC1 cells continued to show increased Alkphos staining when cultured in osteogenic media, while MPC1 cells in growth media showed very little staining (Fig. S1A). In contrast to MPC1 cells, MPC2 cells cultured in growth media displayed strong staining for Alkphos, similar to that of MPC2 cells exposed to osteogenic media (Fig. S1B). Taken together, staining for alizarin red and Alkphos demonstrates that MPC1 and MPC2 cells rapidly differentiate towards the osteogenic lineage, forming mineralized nodules and producing high levels of Alkphos.

### Osteogenic Gene Expression

To quantify expression of specific genes that influence osteogenic differentiation MPC1 and MPC2 cells were cultured in osteogenic media for 0, 7, 14, 21, or 28 days and total RNA was isolated and reverse transcribed into cDNA followed by amplification of genes by qPCR. Comparing MPC1 to MPC2 cells, Sclerostin (*Sost*) expression was significantly higher in MPC2 cells at day 0 (1.4-fold) and day 7 (14.6-fold) (Fig. 2A). No significant differences were observed between cell lines after 14 and 21 days of osteogenic differentiation, whereas MPC2 cells had 86.5% less *Sost* mRNA compared to MPC1 cells at day 28 (Fig. 2A). Compared to day 0, *Sost* expression was significantly increased in MPC1 cells when cultured for 14 (22-fold) and 28 days (7,260-fold) in OM. MPC2 cells had significantly increased *Sost* expression at 7 (35-fold), 14 (31-fold), and 21 days (63-fold) of culture in OM, compared to day 0 (Fig. 2A).

**Figure 2.**
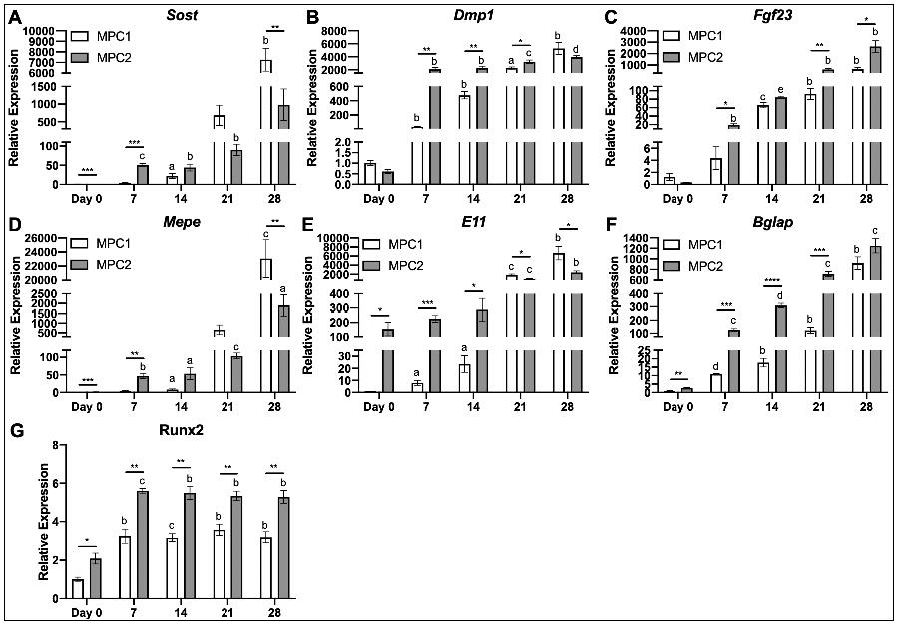
Differentiation of MPC cells abundantly upregulates osteocyte and osteoblast genes. RNA from both MPC1 (white bars) and MPC2 (gray bars) cells were analyzed for several key osteoblast and osteocyte genes over the course of osteogenic differentiation and normalized to β-actin. (A) *Sost* mRNA gradually increased in both cell types over time. MPC2 cells exhibited higher *Sost* mRNA induction at Day 7 compared to MPC1, which normalized at day 14 and 21. By day 28 MPC1 cells expressed significantly more *Sost* mRNA compared to MPC2. Runx2 mRNA levels in undifferentiated cells at Day 0 were modestly elevated in MPC2 cells compared to MPC1. (B) *Dmp1* mRNA was similar at baseline and displayed a dramatic increase at Day 7 of differentiation in both cell types. MPC2 cells maintained elevated Dmp1 levels over MPC1 cells until Day 28. MPC2 cells expressed significantly greater levels of *Dmp1* compared to MPC1 at 7, 14, and 21 days of culture. (C) *Fgf23* mRNA levels between the cell lines were similar at Day 0. *Fgf23* mRNA showed a gradual induction over time with differentiation with both cell types. MPC2 cells maintained elevated *Fgf23* mRNA over MPC1 cells at all time points except for Day 14. (D) *Mepe* mRNA was elevated in MPC2 cells at Day 0 compared to MPC1 cells. With differentiation *Mepe* mRNA significantly rose in MPC2 cells at all time points. For MPC1, *Mepe* expression was only modestly elevated until Day 21. By Day 28 *Mepe* mRNA levels were significantly elevated over MPC2 cells. (E) Compared to MPC1, *E11* mRNA was highly elevated in MPC2 cells at day 0. With differentiation, MPC1 cells showed a gradual increase in *E11* mRNA which surpassed expression levels of MPC2 cells at Day 21 and 28. *E11* mRNA was only significantly elevated vs Day 0 in MPC2 cells after Day 21 and 28 of differentiation. (F) At day 0 *Bglap* mRNA was modestly, but significantly, elevated in MPC2 cells compared to MPC1. With differentiation, *Bglap* mRNA was dramatically upregulated over time in both cell types. Significant differences between MPC1 and MPC2 cells were evident at each time point until Day 28. (G) With differentiation, *Runx2* mRNA significantly increased at day 7 in both cell types compared to day 0. These levels were maintained both throughout differentiation and the elevations between MPC2 cells. (n=3; mean + standard deviation). *p<0.05, **p<0.01, ***p<0.001, and ****p<0.0001 between MPC1 and MPC2 at the time point designated; ^a^p<0.05, ^b^p<0.01, ^c^p<0.001, ^d^p<0.0001, and ^e^p<0.00001 vs day 0 within the same cell line.

Dentin matrix protein-1 mRNA (*Dmp1*) was significantly higher in MPC2 cells compared to MPC1 following 7 (55-fold), 14 (4.8-fold), and 21 days (1.4-fold) of osteogenic differentiation (Fig. 2B). Compared to day 0, MPC1 cells had significantly increased *Dmp1* expression at 7 (37-fold), 14 (469-fold), 21 (2,237-fold), and 28 days (5,193-fold) of culture in OM. MPC2 production of *Dmp1* was significantly increased at 7 (3,412-fold), 14 (3,739-fold), 21 (5,297-fold), and 28 days (6,436-fold) compared to 0 days of OM culture (Fig. 2B).

The expression of fibroblast growth factor-23 (*Fgf23*) mRNA, an endocrine phosphaturic hormone produced by osteocytes, was significantly higher in MPC2 cells compared to MPC1 at 7 (4-fold), 21 (6.7-fold), and 28 days (4-fold) of culture (Fig. 2C). Compared to day 0, *Fgf23* mRNA production was significantly increased in MPC1 cells at 14 (52-fold), 21 (73-fold), and 28 days (511-fold). MPC2 cells had significantly increased Fgf23 expression compared to day 0 when cultured in OM for 7 (54-fold), 14 (238-fold), 21 (1,750-fold), and 28 days (7,429-fold) (Fig. 2C).

Another osteoblast/osteocyte marker, extracellular phosphoglycoprotein, encoded by the *Mepe* gene, was significantly increased in MPC2 cells compared to MPC1 at days 0 (1.4-fold) and 7 (13.6-fold). After 28 days of culture in OM media *Mepe* expression in MPC1 cells was 12-fold higher than MPC2 cells (Fig. 2D). Compared to day 0, *Mepe* mRNA was significantly increased in MPC1 cells with 14 (8-fold) and 28 days (23,052-fold) of osteogenic differentiation. MPC2 cells expressed significantly increased *Mepe*, compared to day 0, at 7 (32-fold), 14 (37-fold), 21 (73-fold), and 28 days (1,334-fold) of osteogenic differentiation (Fig. 2D).

The transmembrane glycoprotein Podoplanin (*E11*) is highly expressed in early stages of osteocyte formation[14]. Expression of *E11* was significantly higher in MPC2 cells compared to MPC1 at day 0 (153-fold), 7 (114-fold) and 14 (12-fold). MPC1 cells produced significantly higher *E11* levels at 21 (0.55-fold) and 28 days (2.7-fold) of osteogenic culture (Fig. 2E). Compared to day 0, MPC1 cells produced significantly increased levels of *E11* at 7 (2-fold), 14 (23-fold), 21 (1,849-fold), and 28 days (6,644-fold) of culture. Osteogenic differentiation of MPC2 cells produced significantly increased *E11* at days 21 (6.7-fold) and 28 (15.7-fold) compared to day 0 (Fig. 2E).

Osteocalcin (*Bglap*) mRNA expression was significantly increased in MPC2 cells compared to MPC1 at day 0 (2.6-fold), day 7 (11.6-fold), day 14 (17.4-fold), and day 21 (5.8-fold). Compared to day 0, MPC1 cells had significantly higher levels of *Bglap* at 7 (10.7-fold), 14 (17.4-fold), 21 (120-fold), and 28 days (905-fold) of osteogenic differentiation (Fig. 2F). For MPC2 cells, *Bglap* expression was significantly increased after 7 (47-fold), 14 (114-fold), 21 (264-fold), and 28 days (463-fold) of exposure to OM (Fig. 2F).

Runt-related transcription factor 2 (*Runx2*) is a master regulator of osteogenic differentiation[15]. Compared to MPC1, MPC2 cells produced significantly greater levels of *Runx2* at every timepoint surveyed, with the greatest difference being 2-fold (Fig. 2G). In MPC1 cells OM induced significant increases in *Runx2* at each time point, all of which were increased by approximately 3-fold compared to day 0. MPC2 cells also expressed significantly more *Runx2* after addition of OM with production increasing about 2.6-fold at every time point measured compared to day 0 (Fig. 2G).

### Effects of PTH and 1,25D Treatment

Bone is acutely sensitive to hormonal signals, many of which directly influence lineage commitment of mesenchymal progenitors. Parathyroid hormone suppresses *Sost*/sclerostin expression both *in vivo* [16] and *in vitro* [9], whereas 1,25D has been shown to influence expression of several osteogenic genes, including *Fgf23* [17]. To assess the ability of PTH and 1,25D to regulate expression of osteogenic genes in MPC cells, MPC1 and MPC2 cells were exposed to differentiation media for 28 days followed by treatment with PTH (50 nM) or 1,25D (10 nM). PTH significantly suppressed *Sost* mRNA in both MPC1 (99.9%) and MPC2 cells (98.8%) (Fig. 3A). Treatment with 1,25D did not alter *Sost* mRNA expression in either cell line. Treatment with PTH virtually abolished *Dmp1* mRNA in both MPC1 (99.2%) and MPC2 (99.4%) cells, whereas exposure to 1,25D resulted in significantly increased production of *Dmp1* mRNA by 2.7-fold in MPC1 and 3.7-fold in MPC2 cells (Fig. 3A). In MPC1 cells, treatment with PTH significantly decreased *Fgf23* mRNA expression by 39%, while no changes were observed in MPC2 cells. Treatment with 1,25D significantly increased *Fgf23* transcripts in both MPC1 (19.8-fold) and MPC2 (3.9-fold) cells. As previous studies have demonstrated that expression of *Sost* and *Dmp1* is suppressed by PTH, and 1,25D induces both *Dmp1* and *Fgf23* expression, our results demonstrate that osteogenic differentiation of MPC cells recapitulates the hormonal responses of bone.

**Figure 3.**
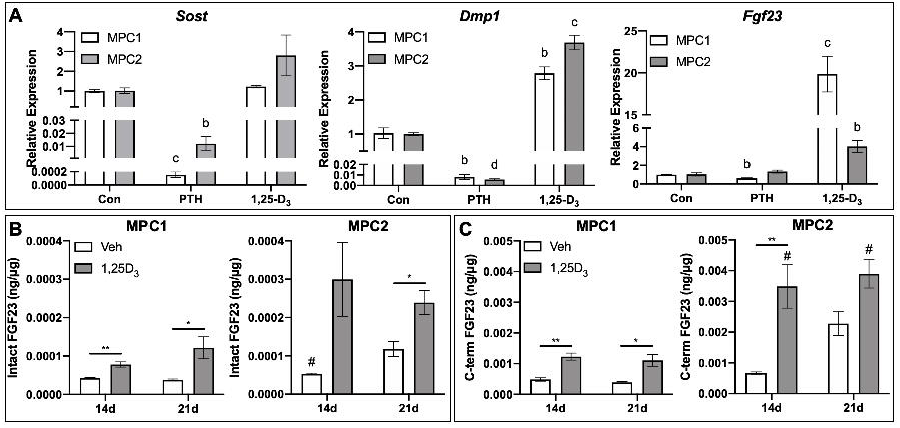
Osteogenic MPC1 and MPC2 cells respond to common endocrine factors. (A) MPC1 (white bars) and MPC2 cells (gray bars) were differentiated for 28 days in osteogenic media and subsequently treated with PTH (50 mM), 1,25D (10 nM), or vehicle control (Con) for 48 hours. *Sost* mRNA was significantly suppressed in both cell types with PTH. *Dmp1* mRNA levels decreased with PTH and increased with 1,25D. *Fgf23* mRNA also decreased with PTH and increased with 1,25D approximately equally with both cell lines. Owing to the induction of *Fgf23* mRNA, MPC1 and MPC2 lines were differentiated for 14 or 21 days then exposed to 1,25D (10^−8^ M, white bars) or vehicle (DMSO, gray bars) for 24 hours to quantify secreted FGF23 protein. (B) In MPC1 cells, iFGF23 increased with 1,25D treatment at both 14 and 21 days of differentiation. In MPC2 cells, 1,25D upregulated FGF23 secretion in the media in 21-day cultures but not at 14 days. (C) Total or cFGF23 was significantly elevated with 1,25D treatment in MPC1 cells after 14 and 21 days of osteogenic differentiation. In MPC2 cells, 1,25D increased cFGF23 release at 21, but not 14 days. MPC2 cells differentiated for 14 or 21 days had higher cFGF23 secretion compared to MPC1 cells at the same timepoints. *p<0.05, **p<0.01, between Veh and 1,25D at the time point designated; ^a^p<0.05, ^b^p<0.01, ^c^p<0.001, and ^d^p<0.0001 compare PTH or 1,25D treatment to Con treatment within the same cell line; #p<0.05 comparing MPC1 and MPC2 at the same treatment and timepoint.

### MPC Cells Secrete FGF23 Protein Regulated by 1,25D

While several cell lines produce *Fgf23* mRNA, few have demonstrated the ability to secrete functional FGF23 protein. As such, we sought to determine not only if MPC cells produced FGF23 protein, but to quantify the extent to which both intact, bioactive FGF23 (iFGF23) and *C*-terminal forms of secreted FGF23 (cFGF23 or “total FGF23” measure both intact and *C*-terminal FGF23 proteolytic fragments) were regulated by 1,25D using two distinct ELISAs. After 14 days of osteogenic differentiation, at baseline (no treatment), MPC1 cells released 4.24×10^−5^ ng/μg of intact FGF23, as measured by ELISA of the cell media, which was normalized to total cell protein content. MPC2 cells produced 5.27×10^−5^ ng/μg, which was significantly increased compared to MPC1 cells, consistent with higher *Fgf23* mRNA content. No significant differences in intact FGF23 were seen between 14 and 21 days of osteogenic differentiation (Fig. 3B). In MPC1 cells 1,25D treatment significantly increased secretion of iFGF23 at both 14 (1.8-fold) and 21 days (3.2-fold) of osteogenic differentiation (Fig. 3B). Exposure to 1,25D increased iFGF23 production at 14 days in MPC2 cells by 5.7-fold; however, this change was not significant. A 2-fold increase in intact FGF23 secretion was observed with 1,25D exposure in MPC2 cells after 21 days of osteogenic culture. Overall, we found that both cell lines secrete more cFGF23 than iFGF23. Secretion of cFGF23 was increased at 14 (2.5-fold) and 21 days (2.8-fold) of culture in MPC1 cells treated with 1,25D. Treatment with 1,25D resulted in a significant increase in cFGF23 of MPC2 cells at 14 days, but not at 21 days. While no differences were observed in cFGF23 production between MPC1 and MPC2 cells in the untreated condition, MPC2 cells exposed to 1,25D secreted significantly more cFGF23 compared to treated MPC1 cells at 14 (2.8-fold) and 21 days (3.5-fold) of culture (Fig. 3C), also consistent with higher *Fgf23* mRNA levels in this line. These data demonstrate that MPC1 and MPC2 cells produce and secrete iFGF23 and cFGF23 protein that are sensitive to 1,25D.

### Adipogenic Differentiation of MPC Cells

Cells of the mesenchymal lineage have the capacity to differentiate into several cell types. As allocation of MSCs towards the osteogenic and adipogenic lineages are inversely proportional[18], the ability of a progenitor cell line to differentiate towards both the adipogenic and osteogenic lineages would present as a useful tool. To determine the ability of MPC cells to differentiate into adipocytes, MPC1 and MPC2 cells were cultured for 4 days in adipogenic media or growth media (as described above). Adipogenic differentiation of both MPC1 (Fig. 4A) and MPC2 (Fig. 4B) cells resulted in increased oil-red-O staining compared to those cultured in growth media.

**Figure 4.**
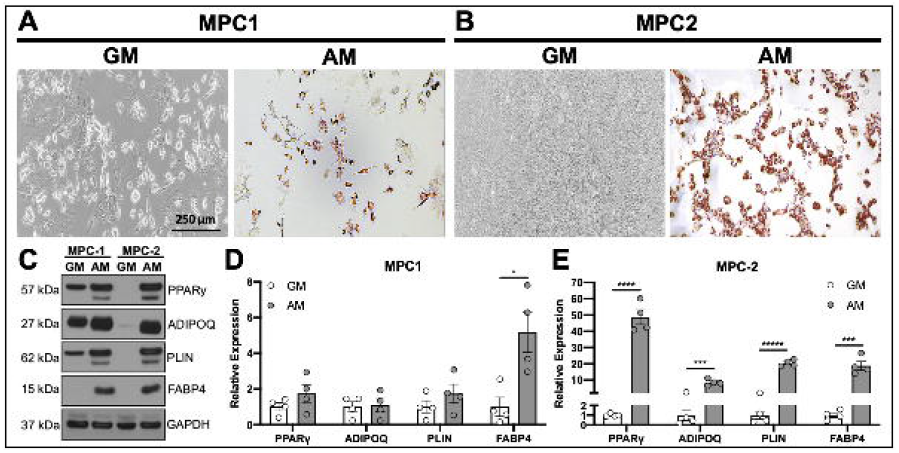
MPC cells undergo robust adipocyte differentiation. MPC cells were cultured for 4 days in growth media (GM) or adipogenic media (AM). Cells were stained with Oil Red O to examine lipid accumulation. (A) MPC1 cells grown in AM stained for Oil Red O while cultures in GM had no staining (bar = 250 μm). (B) MPC2 cells again showed no evidence of lipid formation in GM, whereas nearly all of the cells in view were stained with Oil Red O when exposed to AM. Staining was MPC2 cells was considerably greater than that of MPC1. (C) Protein lysates were assessed from each cell line and media condition for adipogenic proteins PPARγ, ADIPOQ, PLIN and FABP4 by Western blotting. (D) Lysates from four biological replicate experiments were separated by Western blotting and normalized to GAPDH. MPC1 cells only demonstrated a significant increase in FABP4 in AM conditions (gray bars) compared to GM (white bars). Significant increases in PPARγ, ADIPOQ, PLIN and FABP4 under AM conditions were observed in MPC2 cells. *p<0.05, ***p<0.001, ****p<0.0001, *****p<0.00001 vs GM.

To quantify the changes in adipogenic differentiation, MPC cells were exposed to growth media or adipogenic media for 4 days at which time cells were lysed and proteins separated by SDS PAGE for Western blotting (Fig. 4C). Expression of proteins that control adipogenic differentiation, or are a by-product thereof were quantified. Adipogenic differentiation of MPC1 cells did not result in significant increases in peroxisome proliferator-activated receptor gamma (PPARγ), adiponectin (ADIPOQ), or perilipin (PLIN); however, production of fatty acid binding protein 4 (FABP4) was significantly increased by 5-fold (Fig. 4D). In contrast to MPC1 cells, MPC2 cells cultured in adipogenic media had significantly increased expression of adipogenic differentiation markers including PPARγ (48.7-fold), ADIPOQ (8.5-fold), PLIN (20.3-fold), and FABP4 (18.7-fold) as shown by densitometry quantification of Western blots (Figs 5D & E). These data demonstrate that MPC cells are capable of entering into the adipogenic lineage, with MPC2 cells demonstrating greater expression of adipogenic proteins compared to growth media than MPC1 cells.

**Figure 5.**
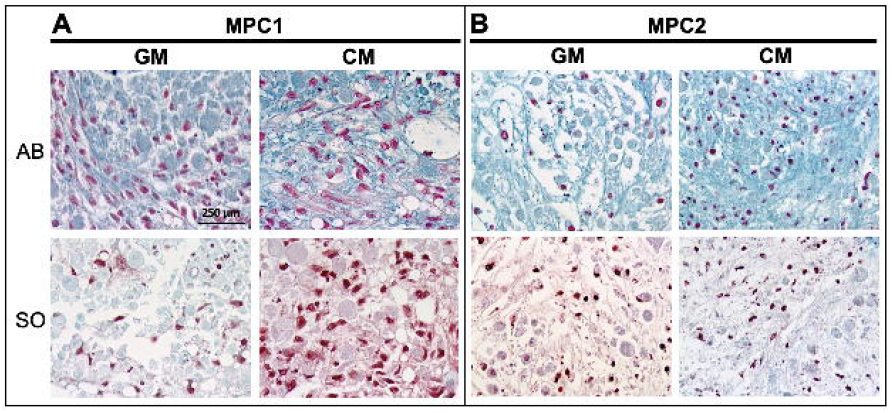
MPC1 and MPC2 cells display chondrogenic differentiation. MPC cells were grown in pellets in either growth media (GM) or chondrogenic media (CM) for 28 days then stained for Alcian Blue (AB) and Safranin-O (SO). MPC1 cells show more robust Safranin O stain under chondrogenic conditions, whereas MPC2 cells displayed strong Alcian Blue staining in chondrogenic conditions, but no clear differences in Safranin-O. (10X; bar = 250 μm)

### Chondrogenic Differentiation of MPC Cells

Studies performed using primary chondrocytes are challenging as these cells are limited by their relatively short life span and the laborious nature of procuring primary cells. Immortalized cells with the capacity to differentiate into chondrocytes provide a consistent source of cells, capable of yielding reproducible results, thus decreasing the need for primary cell isolations. As such, MPC cells were cultured in chondrogenic differentiation media to examine the ability of these cells to enter into the chondrogenic lineage. MPC1 and MPC2 cells were pelleted and exposed to growth media or chondrogenic media, as described above. Pellets were embedded and sectioned followed by staining with Alcian blue and safranin-O. In MPC1 cells, Alcian blue staining was slightly more abundant when cells were cultured in chondrogenic media, but not drastically different (Fig. 5A). Safranin-O staining of MPC1 cells cultured in chondrogenic media was considerably greater than that of cells cultured in growth media (Fig. 5A). MPC2 cells grown in chondrogenic media displayed increased staining of Alcian blue, but no distinguishable differences in safranin-O between growth and chondrogenic culture conditions (Fig. 5B).

To examine growth under chondrogenic conditions, MPC cells were cultured in pellets as described, followed by quantification of the pellet size (Fig. 6A). Compared to growth media, MPC1 cells grown in chondrogenic media displayed a 2.1-fold increase in pellet size (Fig. 6B). MPC2 cells cultured in chondrogenic media displayed no change in pellet size compared to those grown in growth media.

**Figure 6.**
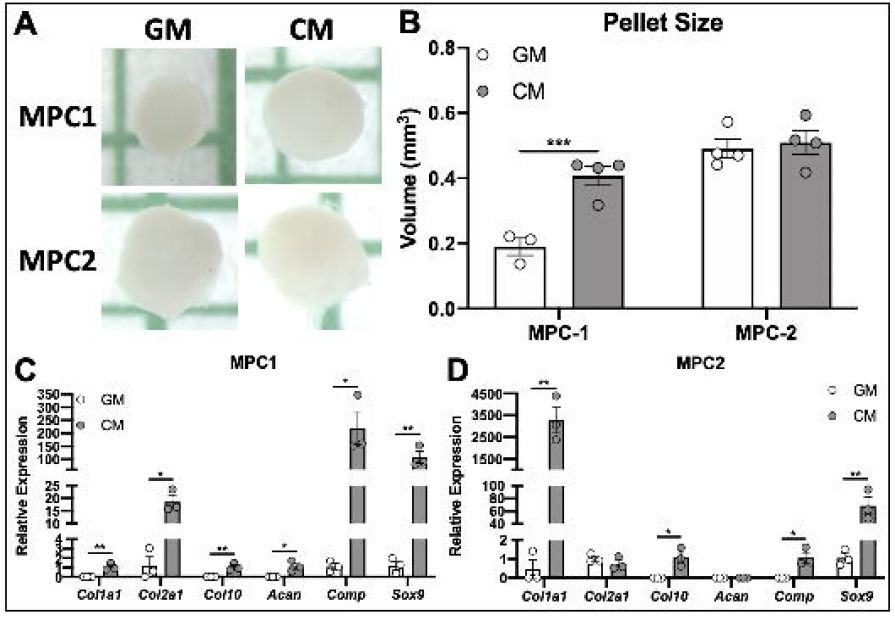
MPC cells grow into cartilaginous pellets and expression chondrogenic genes. MPC cells were cultured in growth media (GM) or chondrogenic media (CM) for 28 days. (A) Under low adherence culture conditions spheroid pellets developed and pellet volumes were measured. MPC1 cells showed increased pellet volume in chondrogenic conditions (gray bars) compared to growth media (white bars). There was no change in pellet volume for MPC2 cells. (B) RNA was analyzed after 28 days of differentiation. For MPC1 cells, chondrogenic conditions (CM) significantly enhanced mRNA levels of chondrocyte markers *Col1a1*, *Col2a1*, *Col10*, *Acan*, *Comp*, and *Sox9*. MPC2 cells differentiated with CM showed robust upregulation of *Col1a1* and *Sox9*. *Col10* and *Comp* mRNA increase was more modest and there was no change in *Col2a1* and *Acan*. n=3; *p<0.05, **p<0.01, ***p<0.001.

To determine if exposure to chondrogenic media induced expression of genes associated with chondrogenesis, MPC cells were cultured in pellets followed by mRNA isolation and qPCR analysis. In MPC1 cells expression of collagen 1a1 (*Col1a1*), collagen 2a1 (*Col2a1*) (15.3-fold), collagen 10 (*Col10*), aggrecan (*Acan*), cartilage oligomeric matrix protein (*Comp*) (200-fold), and SRY-box transcription factor 9 (*Sox9*) (89-fold) were all significantly increased when cultured in chondrogenic media (Fig. 6C). Levels of *Col1a1*, *Col10*, and *Acan* were undetectable in samples grown in growth media, thus fold-change could not be calculated for those samples. In MPC2 cells, mRNA expression of *Col1a1* (6,912-fold), *Col10*, *Comp*, and *Sox9* (65.4-fold) were significantly increased when cultured in chondrogenic media, compared to maintenance growth media. Levels of *Col10*, *Acan,* and *Comp* mRNAs were undetectable in samples grown in growth media. No changes in expression of *Col2a1* or *Acan* were observed between growth and chondrogenic media (Fig. 6D).

Cartilaginous pellets formed by MPC1 and MPC2 cells were embedded and sectioned for immunostaining to examine expression of proteins associated with cartilage development. MPC1 cells exposed to chondrogenic media had strongly increased expression of aggrecan (ACAN) (Fig. 7A). No apparent differences were observed in expression of collagen 1 (COL1) or collagen X (COLX) in MPC1 cells grown in chondrogenic media. Similar to MPC1 cells, chondrogenic differentiation of MPC2 cells resulted in increased expression of ACAN (Fig. 7B). While no differences in COL1 expression were observed when MPC2 cells were exposed to chondrogenic media, immunostaining of COLX was increased in cells grown under chondrogenic conditions compared to growth media (Fig. 7B). Taken together, these data demonstrate that MPC cells are capable of developing into cartilage-like structures that exhibit proteoglycan staining, expression of chondrogenic genes, and production of protein characteristic of cartilage.

**Figure 7.**
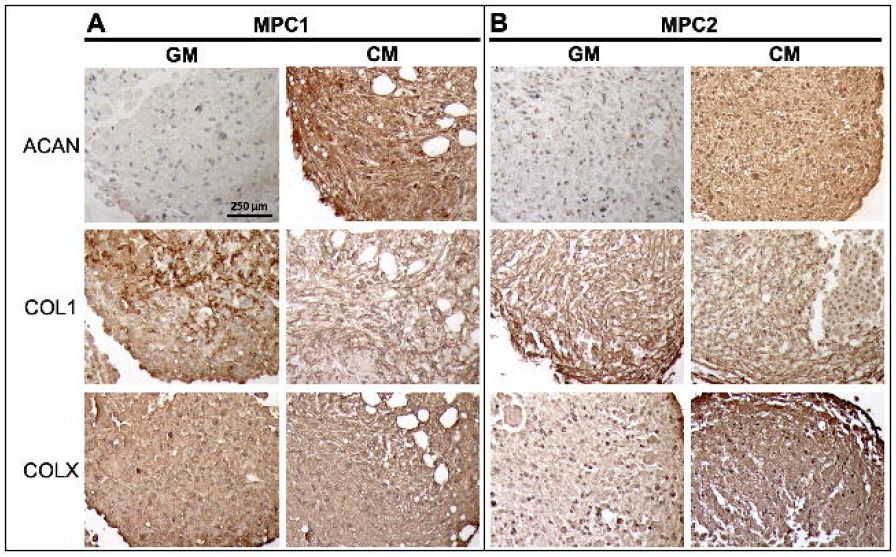
MPC chondrogenic pellets express cartilaginous proteins. MPC cell lines were grown into spheroids within 28 days of culture in growth media (GM) or chondrogenic media (CM). Pellets were sectioned and stained for Aggrecan (ACAN), Type 1 Collagen (COL1) and Type 10 Collagen (COLX). (A) Pellets from MPC1 cells had increased ACAN staining when cultured in CM. No apparent changes in COL1 or COLX were observed in MPC1 cells. (B) MPC2 cells had increased ACAN staining, but less dramatic than MPC1. MPC2 cells showed no change in COL1 but had enhanced COLX staining. (10X; bar = 250 μm)

In sum, we have developed two novel mesenchymal progenitor cell lines that mimic the *in vivo* ability to differentiate into osteoblasts/osteocytes, adipocytes, and chondrocytes. With the capability of responding to exogenous stimuli, known to affect the responses of differentiated cell types, these cells provide a powerful tool to examine responses of progenitors and the differentiated cells. Further, these cells are readily transfected in the undifferentiated state, as we have shown by transfecting GFP (Fig. S2), enabling gene targeting approaches for mechanisms-based experiments within differentiated cell types.

## Discussion

Mesenchymal stem cells serve as the progenitors for multiple tissue types, including adipose, bone, and cartilage. A variety of hormonal, chemical, and physical cues influence lineage allocation of MSCs; however, studying these individual effects *in vivo* present numerous challenges. Despite the availability of several methods for the isolation and characterization of MSCs, obtaining homogenous cultures capable of faithfully replicating bone, cartilage, and fat cell phenotypes remains challenging. Herein, we describe the development of two novel progenitor cell lines, MPC1 and MPC2, which serve as a useful tool to study the properties of mesenchymal progenitors, but also recapitulate the morphological, genetic, and signaling properties of bone, cartilage, and adipose. These cells can be routinely used to study the factors regulating the transition between these lineages, as well as the terminally differentiated cell types produced from differentiation.

Immortalization of MPC1 and MPC2 cells was accomplished by the expression of the temperature sensitive large T-antigen, enabling proliferation at 33°C and differentiation at 37°C. The ability to restrict proliferation is a useful feature for the study of some cell types. In particular, osteocytes, which are the terminally differentiated form of osteoblasts, are not proliferative *in vivo*[19]. Thus, osteocyte cell lines that maintain proliferative capacity may not fully replicate the osteocyte phenotype. Incorporation of the temperature sensitive large T-antigen into these cells enables the ability to restrict proliferation at the time point of choosing[20], which may be equally useful for the study of hypertrophic chondrocytes.

In previous work, our group showed that MSCs isolated from mouse bone marrow were capable of differentiating into mineralized nodules[9]. These stem cell-derived osteocyte (SCD-O) cultures formed bone-like nodules containing osteocytes which replicated the morphology and structure of osteocytes in cortical bone. As osteocyte responses rely heavily on connections to the matrix, this three-dimensional environment is essential for the replication of the *in vivo* condition. When primary osteocytes are isolated, they are removed from the native matrix environment which defines their morphology and regulates contextual behavior. Thus, primary osteocytes grown in culture dishes quickly lose their osteocyte-like features.

Alizarin red and alkaline phosphatase staining of MPC cells demonstrated formation of mineralized bone-like nodules, which were evident as early as 7-14 days. Thus, when grown in osteogenic conditions MPC cultures mimic the *in vivo* matrix environment. Some of the earlier produced osteocytic cell lines, such as MLO-Y4 [21] and MLO-A5 [22] are cultured on a monolayer and thus are also devoid of the matrix environment that is essential for osteocyte responses. However, there have been several recent cell lines produced that form mineralized bone-like structures. These include the IDG-SW3 [20], OCY-454 [23], and OmGFP66 [24]. In particular, after 28 days of culture in osteogenic media OmGFP66 cells formed highly organized 3D bone-like structures that resembled trabeculae. These cultures had well-defined osteocyte spacing with the characteristic osteocyte morphology [24]. Importantly, MPC1 and MPC2 cells form mineralized nodules rapidly in culture, providing the matrix environment necessary to support osteocyte differentiation and production/secretion of osteocyte products. In contrast to other commonly used osteocytic cell lines, MPC cells did not require collagen coating of culture dishes to induce or maintain differentiation.

In addition to the formation of abundant matrix, MPC cells also produced all of the characteristic markers of osteocytes including the early osteocyte marker E11. Expression of genes that are produced in mature osteocytes, including *Sost*, *Dmp1*, *Fgf23*, and *Mepe* were also extremely abundant. Notably, osteogenic differentiation resulted in production of these osteocyte-specific genes on the order of 2,000 – 22,000-fold higher than in the undifferentiated stage. Production of these genes at such high levels not only highlights the ability of MPC cultures to serve as a unique source of osteocytic cells, but also serves as an excellent model to study the transcriptional regulation of each of these genes. Sclerostin in particular has received great attention in the last decade as it is an inhibitor of Wnt/Lrp signaling, and the target of the neutralizing monoclonal Ab romosozumab [25], a drug recently approved to treat osteoporosis. Our data show that MPC cells treated with PTH have a dramatic reduction of *Sost* expression as well as *Dmp1*. These results align well with previously published *in vivo* and *in vitro* studies demonstrating hormonal control of these genes and that MPC cells replicate key functional responses of osteocytic genes.

The osteocyte-produced hormone FGF23 has been studied in a number of cell models. The MPC cell lines provide a novel system for testing FGF23 biology in regard to this hormone’s expression and actions. MPC cells express FGF23 in response to 1,25D, similar to osteosarcoma lines such as the rat osteoblast/osteocyte cell line UMR-106 [26,27]. The UMR-106 line also increased *Fgf23* mRNA when challenged with hypoxia-mimetics [28] and iron chelation [29]. Further, response elements in *Fgf23* promoter fragments were activated in response to 1,25D in human K562 leukemia cells [30]. The human osteoblastic cell line U2OS also properly expressed *Fgf23* cDNA, and gene targeting was used to identify key glycosylation/phosphorylation events necessary for FGF23 production [31]. The disadvantage of these lines is that they do not appear to express endogenous FGF23 protein at readily quantifiable levels.

The mouse osteoblast-like cell line MC3T3-E1 has also been used to study FGF23 expression and its possible roles in osteoblast function [32], as well as its promoter elements [33]. Similar to UMR-106 and U2OS cells; however, the detection of measurable FGF23 protein has been elusive in these cells. Whether the lack of mature protein has to do with portions of the protein folding/secretory pathways down regulated over time in the cells or the fact that FGF23 can be proteolytically inactivated prior to secretion [29,34] remains unclear. When differentiated, the MPC cell lines expressed both FGF23 mRNA, and secreted protein as detected by both ‘Intact’ and ‘*C*-terminal’ FGF23 ELISAs. This capability allows the future study of FGF23 protein processing using inhibitors/activators, as well as the mechanisms underlying secondary modification of the mature FGF23 protein. Primary cultures of isolated osteoblasts from rodents also express *Fgf23* mRNA [35]; however, these cells are somewhat limited in that, as for primary differentiated cells, they are best transfected using specialized viral expression vectors. Groups have also used differentiated primary bone marrow stromal cells (BMSC) to examine FGF23 production under hypoxic conditions and in response to pro-inflammatory stimuli [33]. Like primary osteoblasts/osteocytes, these cells are typically transduced with viral vectors, and can therefore not be easily targeted by standard genomic modifying reagents such as CRISPR.

The MPC lines have the distinct advantage that genes can be targeted by standard transfection techniques during the undifferentiated growth phase, then tested for cell function in the differentiated osteocyte-like state or as we show, during active mineralization. More recently, a clonal osteogenic cell line, OmGFP66, was developed by immortalization of primary bone cells from mice expressing a membrane-targeted GFP driven by the Dmp1-promoter [24]. This line increased FGF23 expression upon differentiation, similar to the MPC lines. FGF23 was also studied in a model osteocyte-like cell line, IDG-SW3, which demonstrated that ^35^S-labeled FGF23 was cleaved to smaller fragments which were constitutively secreted [36]. In contrast, intact bioactive FGF23 was more efficiently stored in differentiated than in undifferentiated IDG-SW3 cells. Following osteogenic differentiation of IDG-SW3 cultures, basal *Fgf23* mRNA was dose-dependently up-regulated by pro-inflammatory cytokines TNF, IL-1β and TWEAK, and bacterial LPS [37]. cFGF23 and iFGF23 protein levels also increased, but intact protein only in the presence of furin inhibitors, supporting that FGF23 cleavage controls this hormone’s bioavailability. Further, some osteocyte cell lines, such as MLO-Y4 express negligible levels of FGF23 in the basal state; however, modest induction can be seen for *Fgf23* mRNA and other osteocyte genes with culture under 3D conditions. Cell lines of other lineages, including HK2 (Human Kidney-2, proximal tubule-like) cells have been tested, and *Fgf23* and osteopontin mRNAs were expressed in these cells when incubated with TGFβ1; however, these levels were not altered in HK2 cells when treated with 1,25D and high phosphate levels [38]. Although FGF23 is not normally expressed to a significant degree in kidney *in vivo*, this cell model may be useful for testing FGF23 expression under specific pathologic conditions. Thus, the novel MPC cell lines faithfully recapitulate many of the critical features of osteocytes, and will allow further understanding of FGF23 transcription, protein modification, and secretion.

While there are several cell lines capable of recreating osteoblast and osteocyte phenotypes, MPC cells provide a distinct advantage with the capability to also differentiate into the adipogenic and chondrogenic lineages. MPC1 and MPC2 cells readily formed adipocytes after only 5 days of exposure to adipogenic media, much faster than the 2-3 weeks of adipogenic differentiation required of other MSC lines [5,6]. MPC2 cells responded to adipogenic conditions with greater increases in adipogenic proteins compared to MPC1 cells. MPC2 cells consistently produced high levels of the transcriptional regulator of adipogenesis, PPARγ as well as other proteins characteristic of adipocytes including adiponectin, perilipin, and fatty acid binding protein 4. Recent studies have shown that accumulation of adipose tissue within the bone marrow is detrimental to skeletal health and is associated with aging and disuse [39,40]. Additionally, use of certain drugs, such as rosiglitazone increase bone marrow adiposity, but these effects can be attenuated by exercise [41]. Several studies have demonstrated that the mechanical contribution of exercise restrict entry of mesenchymal progenitors into the adipogenic lineage [42,43], effects that are influence by actin cytoskeletal organization [10,44,45], and even shuttling of actin monomers to into the nucleus to regulate YAP/TAZ dynamics [46]. As the balance between osteogenic and adipogenic fate has important implications for health and disease, MPC cells are a good model to study the genetic and phenotypic changes governing the balance of these two mesenchymal tissues.

Several different culture models have been developed to study chondrocytes *in vitro* including explant models, three-dimensional culture systems, and cells grown in monolayers [1]. When grown on a monolayer, devoid of the surrounding matrix, primary chondrocytes readily undergo de-differentiation with progressive loss of collagen type II and aggrecan [47,48]. Culturing chondrocytes within conditions enabling a rounded morphology promotes maintenance of the chondrocyte phenotype [49,50], as such we allowed MPC cultures to grow within pellets. MPC cells grew into cartilaginous pellets which stained for both alcian blue and safranin-O. MPC1 cells demonstrated increased growth capacity compared to MPC2 cells, and more consistently expressed chondrocyte markers. MPC1 cells exposed to chondrogenic media produced collagens 1, 2, and 10, as well as the chondrocyte transcription factor *Sox9* and the matrix molecules aggrecan and *Comp*. MPC1 cells also stained strongly for aggrecan protein. As such, MPC1 cells reproduced the cartilage phenotype more readily than MPC2 cells. Other cell lines including CFK2, which was established from fetal rat calvariae [51], and ATDC5, an embryonal carcinoma cell line isolated from a differentiating culture of AT805 teratocarcinoma [52], reproduce the cartilage phenotype *in vitro*. With the ability to produce cartilage-specific genes and grow within a 3D environment, MPC cells represent a novel cell line to study chondrocyte development, particularly the progression from the progenitor stage.

While both MPC1 and MPC2 cell lines have several beneficial features, there remain limitations in the use of these cells. As MPC cells are derived from single cell clones, they are clonally similar in the undifferentiated state. This characteristic is beneficial to achieve reproducible results; however, studies seeking to replicate the heterogenous population of the bone marrow niche may opt for a culture system such as the SCD-O [9] which are not clonal nor immortalized. The immortalized nature of these cells may also be a limitation for some studies, as the process of introducing the immortalization vector may alter responses. We’ve attempted to circumvent this by using the temperature sensitive large T-antigen, thus the proliferative capacity of the cells can be modulated as needed. Despite some of these limitations, MPC cells produce very high levels of osteogenic genes, especially *Sost* and *Fgf23*. Few cell lines secrete FGF23 protein, in particular intact FGF23, a feature that is asset of these cells. In addition to producing these factors, the ability to MPC cells to differentiate into multiple lineages allows investigators the opportunity to study the transition between mesenchymal cell types. An additional strength of this study was that differentiation of MPC cells was achieved across 5 different laboratories. In particular, osteogenic differentiation results were consistent when performed at the University of Adelaide and at Indiana University. The consistent growth and differentiation of these cells among several investigators enables greater confidence in achieving reproducible results, providing increased rigor in future studies.

In summary, we generated and characterized two novel multi-potent cell lines useful for the study of the undifferentiated mesenchymal progenitor state as well as the differentiated osteoblast/osteocyte, chondrocyte, and adipocyte lineages. When cultured in osteogenic conditions, these cells produce abundant mineralized matrix and are capable of expressing the full profile of genes from the MSC precursor, to osteoblast, to osteocytes. MPC cells also quickly differentiate into fat cells, providing a much faster culture model than many currently available cell lines. These cells provide a novel tool to study factors that regulate MSC differentiation as well as the differentiated state of these mesenchymal cell types.

---

**Figure S1: Osteogenic differentiation and alkaline phosphatase staining of MPC cells.** MPC cells were cultured in growth media (GM) or osteogenic media (OM) and stained for alkaline phosphatase. (A) In MPC1 cells Alkphos staining began to appear in cultures grown in osteogenic media at day 14, with large differences at days 21 and 28 compared to growth media. (B) MPC2 cells displayed strong staining at day 21 in osteogenic media. After 28 days in culture MPC2 cells grown in growth media also had strong Alkphos staining. Each cell line was tested in triplicate. Images were captured using a 10X magnification lens.

**Figure S2: Transfection with GFP.** To establish the ability of MPC cells to be transfected, MPC2 cells were cultured in growth media (GM) at 33°C. A vector containing eGFP was transfected into the cells using Fugene-6 HD. Images were captured using a fluorescent microscope (Leica) 24 h after transfection. (10X; bar = 200 μm)

## References

1 Kartsogiannis V, Ng KW. Cell lines and primary cell cultures in the study of bone cell biology [Review] [in English]. Mol Cell Endocrinol 2004;228(1-2):79–102.

2 Ramakrishnan A, Torok-Storb B, Pillai MM. Primary marrow-derived stromal cells: isolation and manipulation. Methods Mol Biol 2013;1035:75–101.

3 Ullah I, Subbarao RB, Rho GJ. Human mesenchymal stem cells - current trends and future prospective. Biosci Rep 2015;35(2).

4 Galarza Torre A, Shaw JE, Wood A et al. An immortalised mesenchymal stem cell line maintains mechano-responsive behaviour and can be used as a reporter of substrate stiffness. Scientific reports 2018;8(1):8981.

5 Siska EK, Weisman I, Romano J et al. Generation of an immortalized mesenchymal stem cell line producing a secreted biosensor protein for glucose monitoring. PloS one 2017;12(9):e0185498.

6 Aomatsu E, Takahashi N, Sawada S et al. Novel SCRG1/BST1 axis regulates self-renewal, migration, and osteogenic differentiation potential in mesenchymal stem cells. Scientific reports 2014;4:3652.

7 Huang S, Xu L, Sun Y et al. An improved protocol for isolation and culture of mesenchymal stem cells from mouse bone marrow. J Orthop Translat 2015;3(1):26–33.

8 Peister A, Mellad JA, Larson BL et al. Adult stem cells from bone marrow (MSCs) isolated from different strains of inbred mice vary in surface epitopes, rates of proliferation, and differentiation potential. Vol 1032004.

9 Thompson WR, Uzer G, Brobst KE et al. Osteocyte specific responses to soluble and mechanical stimuli in a stem cell derived culture model. Scientific reports 2015;5:11049.

10 Thompson WR, Yen SS, Uzer G et al. LARG GEF and ARHGAP18 orchestrate RhoA activity to control mesenchymal stem cell lineage. Bone 2018;107:172–180.

11 Thompson WR, Keller BV, Davis ML et al. Low-Magnitude, High-Frequency Vibration Fails to Accelerate Ligament Healing but Stimulates Collagen Synthesis in the Achilles Tendon. Orthop J Sports Med 2015;3(5).

12 Cary RL, Waddell S, Racioppi L et al. Inhibition of Ca(2)(+)/calmodulin-dependent protein kinase kinase 2 stimulates osteoblast formation and inhibits osteoclast differentiation. Journal of bone and mineral research: the official journal of the American Society for Bone and Mineral Research 2013;28(7):1599–1610.

13 McCoy SY, Falgowski KA, Srinivasan PP et al. Serum xylosyltransferase 1 level increases during early posttraumatic osteoarthritis in mice with high bone forming potential. Bone 2012;51(2):224–231.

14 Staines KA, Prideaux M, Allen S et al. E11/Podoplanin Protein Stabilization Through Inhibition of the Proteasome Promotes Osteocyte Differentiation in Murine in Vitro Models. J Cell Physiol 2016;231(6):1392–1404.

15 Komori T, Yagi H, Nomura S et al. Targeted disruption of Cbfa1 results in a complete lack of bone formation owing to maturational arrest of osteoblasts. Cell 1997;89(5):755–764.

16 Bellido T, Ali AA, Gubrij I et al. Chronic elevation of parathyroid hormone in mice reduces expression of sclerostin by osteocytes: a novel mechanism for hormonal control of osteoblastogenesis [in eng]. Endocrinology 2005;146(11):4577–4583.

17 Noonan ML, White KE. FGF23 Synthesis and Activity. Curr Mol Biol Rep 2019;5(1):18–25.

18 Uzer G, Fuchs RK, Rubin J et al. Concise Review: Plasma and Nuclear Membranes Convey Mechanical Information to Regulate Mesenchymal Stem Cell Lineage. Stem cells 2016;34(6):1455–1463.

19 Prideaux M, Findlay DM, Atkins GJ. Osteocytes: The master cells in bone remodelling. Curr Opin Pharmacol 2016;28:24–30.

20 Woo SM, Rosser J, Dusevich V et al. Cell line IDG-SW3 replicates osteoblast-to-late-osteocyte differentiation in vitro and accelerates bone formation in vivo. Journal of bone and mineral research: the official journal of the American Society for Bone and Mineral Research 2011;26(11):2634–2646.

21 Kato Y, Windle JJ, Koop BA et al. Establishment of an Osteocyte-like Cell Line, MLO-Y4. Journal of Bone and Mineral Research 1997;12(12):2014–2023.

22 Kato Y, Boskey A, Spevak L et al. Establishment of an osteoid preosteocyte-like cell MLO-A5 that spontaneously mineralizes in culture. Journal of bone and mineral research: the official journal of the American Society for Bone and Mineral Research 2001;16(9):1622–1633.

23 Spatz JM, Wein MN, Gooi JH et al. The Wnt Inhibitor Sclerostin Is Up-regulated by Mechanical Unloading in Osteocytes in Vitro. The Journal of biological chemistry 2015;290(27):16744–16758.

24 Wang K, Le L, Chun BM et al. A Novel Osteogenic Cell Line That Differentiates Into GFP-Tagged Osteocytes and Forms Mineral With a Bone-Like Lacunocanalicular Structure. Journal of bone and mineral research: the official journal of the American Society for Bone and Mineral Research 2019;34(6):979–995.

25 McClung MR, Grauer A, Boonen S et al. Romosozumab in postmenopausal women with low bone mineral density. N Engl J Med 2014;370(5):412–420.

26 Kolek OI, Hines ER, Jones MD et al. 1alpha,25-Dihydroxyvitamin D3 upregulates FGF23 gene expression in bone: the final link in a renal-gastrointestinal-skeletal axis that controls phosphate transport [in eng]. American journal of physiology Gastrointestinal and liver physiology 2005;289(6):G1036–1042.

27 Farrow EG, Davis SI, Ward LM et al. Molecular analysis of DMP1 mutants causing autosomal recessive hypophosphatemic rickets [in eng]. Bone 2009;44(2):287–294.

28 Hum JM, Clinkenbeard EL, Ip C et al. The metabolic bone disease associated with the Hyp mutation is independent of osteoblastic HIF1alpha expression. Bone reports 2017;6:38–43.

29 Farrow EG, Yu X, Summers LJ et al. Iron deficiency drives an autosomal dominant hypophosphatemic rickets (ADHR) phenotype in fibroblast growth factor-23 (Fgf23) knock-in mice [in eng]. Proc Natl Acad Sci U S A 2011;108(46):E1146–1155.

30 Kaneko I, Saini RK, Griffin KP et al. FGF23 gene regulation by 1,25-dihydroxyvitamin D: opposing effects in adipocytes and osteocytes. J Endocrinol 2015;226(3):155–166.

31 Tagliabracci VS, Engel JL, Wiley SE et al. Dynamic regulation of FGF23 by Fam20C phosphorylation, GalNAc-T3 glycosylation, and furin proteolysis. Proc Natl Acad Sci U S A 2014;111(15):5520–5525.

32 Shalhoub V, Ward SC, Sun B et al. Fibroblast Growth Factor 23 (FGF23) and Alpha-Klotho Stimulate Osteoblastic MC3T3.E1 Cell Proliferation and Inhibit Mineralization [in Eng]. Calcif Tissue Int 2011.

33 David V, Martin A, Isakova T et al. Inflammation and functional iron deficiency regulate fibroblast growth factor 23 production. Kidney Int 2015;22(6):1020–1032.

34 Benet-Pages A, Lorenz-Depiereux B, Zischka H et al. FGF23 is processed by proprotein convertases but not by PHEX [in eng]. Bone 2004;35(2):455–462.

35 Liu S, Tang W, Fang J et al. Novel regulators of Fgf23 expression and mineralization in Hyp bone [in eng]. Molecular endocrinology 2009;23(9):1505–1518.

36 Yamamoto H, Ramos-Molina B, Lick AN et al. Posttranslational processing of FGF23 in osteocytes during the osteoblast to osteocyte transition. Bone 2016;84:120–130.

37 Ito N, Wijenayaka AR, Prideaux M et al. Regulation of FGF23 expression in IDG-SW3 osteocytes and human bone by pro-inflammatory stimuli. Molecular and cellular endocrinology 2015;399:208–218.

38 Sugiura H, Matsushita A, Futaya M et al. Fibroblast growth factor 23 is upregulated in the kidney in a chronic kidney disease rat model. PLoS One 2018;13(3):e0191706.

39 de Abreu MR, Wesselly M, Chung CB et al. Bone marrow MR imaging findings in disuse osteoporosis. Skeletal Radiol 2011;40(5):571–575.

40 Woods GN, Ewing SK, Sigurdsson S et al. Greater Bone Marrow Adiposity Predicts Bone Loss in Older Women. Journal of bone and mineral research: the official journal of the American Society for Bone and Mineral Research 2020;35(2):326–332.

41 Styner M, Pagnotti GM, Galior K et al. Exercise Regulation of Marrow Fat in the Setting of PPARgamma Agonist Treatment in Female C57BL/6 Mice. Endocrinology 2015;156(8):2753–2761.

42 Sen B, Xie ZH, Case N et al. Mechanical Strain Inhibits Adipogenesis in Mesenchymal Stem Cells by Stimulating a Durable beta-Catenin Signal [Article] [in English]. Endocrinology 2008;149(12):6065–6075.

43 Styner M, Thompson WR, Galior K et al. Bone marrow fat accumulation accelerated by high fat diet is suppressed by exercise. Bone 2014;64(0):39–46.

44 Sen B, Xie Z, Case N et al. mTORC2 regulates mechanically induced cytoskeletal reorganization and lineage selection in marrow-derived mesenchymal stem cells. Journal of bone and mineral research: the official journal of the American Society for Bone and Mineral Research 2014;29(1):78–89.

45 Thompson WR, Guilluy C, Xie Z et al. Mechanically activated Fyn utilizes mTORC2 to regulate RhoA and adipogenesis in mesenchymal stem cells. Stem cells 2013;31(11):2528–2537.

46 Sen B, Xie Z, Uzer G et al. Intranuclear Actin Regulates Osteogenesis. Stem cells 2015;33(10):3065–3076.

47 Takigawa M, Shirai E, Fukuo K et al. Chondrocytes dedifferentiated by serial monolayer culture form cartilage nodules in nude mice. Bone Miner 1987;2(6):449–462.

48 Lefebvre V, Garofalo S, Zhou G et al. Characterization of primary cultures of chondrocytes from type II collagen/beta-galactosidase transgenic mice. Matrix Biol 1994;14(4):329–335.

49 Bonaventure J, Kadhom N, Cohen-Solal L et al. Reexpression of cartilage-specific genes by dedifferentiated human articular chondrocytes cultured in alginate beads. Experimental cell research 1994;212(1):97–104.

50 Hauselmann HJ, Fernandes RJ, Mok SS et al. Phenotypic stability of bovine articular chondrocytes after long-term culture in alginate beads. Journal of cell science 1994;107 (Pt 1):17–27.

51 Bernier SM, Goltzman D. Regulation of expression of the chondrocytic phenotype in a skeletal cell line (CFK2) in vitro. Journal of bone and mineral research: the official journal of the American Society for Bone and Mineral Research 1993;8(4):475–484.

52 Atsumi T, Miwa Y, Kimata K et al. A chondrogenic cell line derived from a differentiating culture of AT805 teratocarcinoma cells. Cell Differ Dev 1990;30(2):109–116.

