## Supplementary figures and images for "Generation of two Multipotent Mesenchymal Progenitor Cell Lines Capable of Osteogenic, Mature Osteocyte, Adipogenic, and Chondrogenic Differentiation"

### Supplementary Data

Figure S1

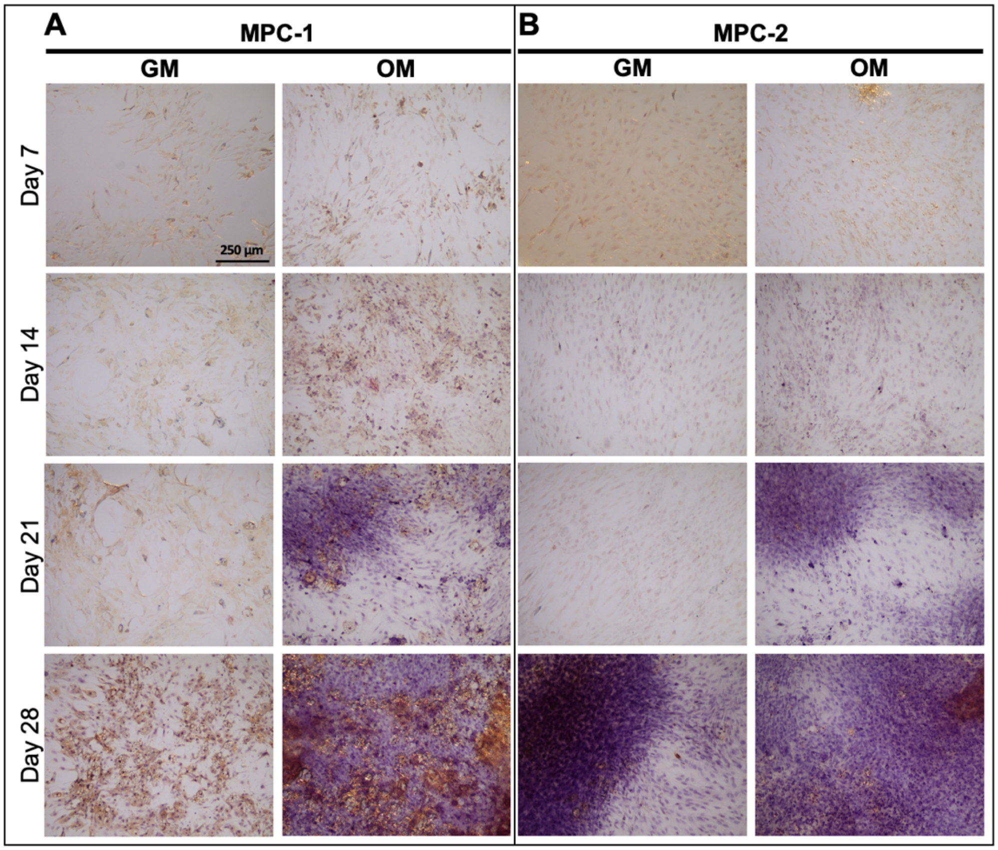


Figure S2

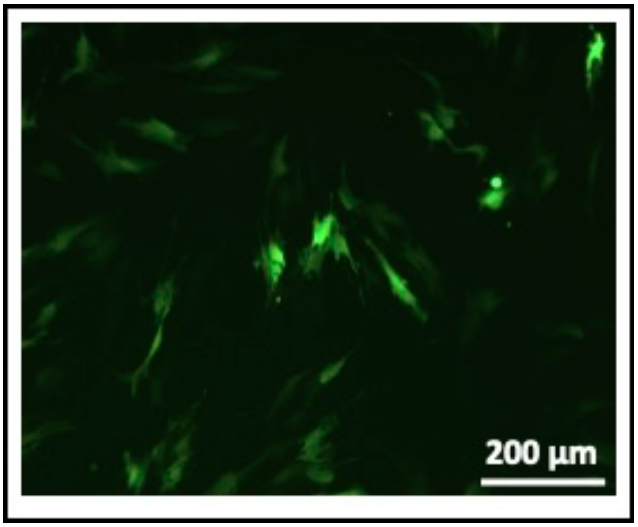
